# Experience dependent plasticity of higher visual cortical areas in the mouse

**DOI:** 10.1101/2023.01.19.524689

**Authors:** Rosie Craddock, Asta Vasalauskaite, Adam Ranson, Frank Sengpiel

## Abstract

Experience dependent plasticity in the visual cortex is a key paradigm for the study of mechanisms underpinning learning and memory. Despite this, studies involving manipulating visual experience have largely been limited to the primary visual cortex, V1, across various species. Here we investigated the effects of monocular deprivation (MD) on the ocular dominance and orientation selectivity of neurons in 4 visual cortical areas in the mouse: the binocular zone of V1 (V1b), the putative ‘ventral stream’ area LM and the putative ‘dorsal stream’ areas AL and PM. We employed two-photon calcium imaging to record neuronal responses in young adult mice before MD, immediately after MD, and following binocular recovery. Ocular dominance shifts following MD were greatest in LM and smallest in AL and PM; in all areas these shifts were mediated primarily through a reduction of deprived-eye responses, and a smaller increase in response through the non-deprived eye. The ocular dominance index recovered to pre-MD levels within 2 weeks in all areas. MD caused a modest reduction in orientation selectivity of deprived-eye responses in V1b, LM and AL, but not PM. Our results suggest that changes in ocular dominance in higher visual areas are not simply inherited from V1.

## Introduction

Since the pioneering work by Hubel and Wiesel on cats and monkeys in the 1960s and 1970s (Wiesel and Hubel, 1963; Hubel et al., 1977) ocular dominance (OD) plasticity has become one of the classic paradigms for the study of experience-dependent brain plasticity. Occluding one eye by monocular deprivation (MD) shifts responses of visual cortical neurons towards the open eye, in particular during a critical period early in life Hubel Wiesel (Hubel and Wiesel, 1970; Olson and Freeman, 1978). It also results in a substantial loss of visual acuity in the deprived eye (Dews and Wiesel, 1970) (Harwerth et al., 1983), a condition known as amblyopia. Studies of OD plasticity have for over 50 years focused on the primary visual cortex (V1) although it is accepted that not all visual deficits observed in amblyopic animals can be accounted for by the abnormalities of responses seen in V1 (see (Kiorpes and Daw, 2018).

Although neurons in the mouse primary visual cortex, unlike most primate and carnivore species, are not organised into ocular dominance columns and are overall more strongly dominated by the contralateral eye (Dräger, 1975), they nevertheless respond to MD with a characteristic OD shift towards the open eye, as originally demonstrated in single-cell recording studies (Dräger, 1978) (Gordon and Stryker, 1996) and later confirmed using two-photon imaging (Mrsic-Flogel et al., 2007). Mouse V1 is most susceptible to the effects of MD during the critical period, peaking at P28 (Gordon and Stryker, 1996), but OD shifts can still be observed in older animals (Sawtell et al., 2003), and in some cases even in animals up to at least 8 months of age (Greifzu et al., 2014). From a number of studies a consensus has emerged that OD plasticity in V1 of young mice within the classical critical period is mechanistically different from that observed in animals that are two months of age or older (for reviews see (Espinosa and Stryker, 2012; Levelt and Hübener, 2012). During the critical period, a brief MD of 3 days results in a Hebbian, LTD-type depression of deprived-eye responses (Bear et al., 1990) while slightly longer MD of 6 days is characterised by a homeostatic upregulation of both the open-eye and (to a lesser extent) the deprived-eye responses which requires tumor necrosis factor-alpha (Kaneko et al., 2008a). Upon re-opening of the deprived eye responses through both eyes return to baseline levels, and this recovery is dependent upon brain-derived neurotrophic factor via tyrosine kinase B receptor activation (Kaneko et al., 2008b). In contrast, OD plasticity in adult mice is apparent only after 6-7 days of MD and is characterised solely by an increase in the open-eye response, with no change in the deprived-eye response, and this is NMDA receptor dependent (Sawtell et al., 2003; Ranson et al., 2012).

Despite the wealth of knowledge concerning OD plasticity in the primary visual cortex of mice very little is known about higher visual areas (HVAs) in this respect. Recent studies have revealed at least nine retinotopically organised HVAs surrounding V1 (Wang and Burkhalter, 2007), and it has been suggested on the basis of axonal projection patterns that they are organised into to two groups which may be equivalent to the dorsal and ventral processing streams known from the primate visual system (Wang et al., 2011; Wang et al., 2012). Despite some anatomical crosstalk between the two streams they appear to be functionally distinct based on their spatiotemporal selectivity (Murakami et al., 2017), although other studies have highlighted some inconsistencies.

Here we explore OD plasticity in higher visual areas that are direct downstream targets of binocular V1, including areas both in the putative dorsal stream (AL, PM) and ventral stream (LM). We specifically aim to address the questions whether the higher visual areas simply inherit plastic changes occurring in V1 or independently express experience-dependent plasticity, and whether dorsal and ventral streams exhibit different forms or degrees of plasticity. We use two-photon calcium imaging to analyse the effects of MD on responses of individual neurons and subsequent recovery under binocular vision.

## Materials and Methods

### Animals

All experimental procedures were carried out in accordance with the UK Animals (Scientific Procedures) Act 1986 and European Commission directive 2010/63/EU. Mice expressing GCaMP6f in excitatory neurons were generated by crossing the Ai95D line (Jax, 024105) with the CaMkII-alpha-cre T29-1 line (Jax, 005359). Experiments were carried out on young adult mice of either gender aged 2-3 months, housed under normal light conditions (12h light, 12h dark). All recordings were made during the light period.

### Surgical preparation

Surgical procedures were in line with recently published recommendations (Barkus et al., 2022). All surgical procedures were conducted under aseptic conditions. Prior to cranial window surgery, animals were administered with the antibiotic Baytril (5mg/kg, s.c.) and the anti-inflammatory drugs Rimadyl (5mg/kg, s.c.) and Dexamethasone (0.15mg/Kg, i.m.). Anaesthesia was induced at a concentration of 4% Isoflurane in oxygen, and maintained at 1.5-2% for the duration of the surgery. Once fully anaesthetised animals were secured in a stereotaxic frame (David Kopf Instruments, Tujunga, CA, USA) and the scalp and periosteum were removed from the dorsal surface of the skull. A custom head plate (see Fig. 1B, (Barkus et al., 2022) was attached to the cranium using dental cement (Super Bond, C&B), with an aperture centred over the right hemisphere visual cortex. A 3- mm circular craniotomy was then made, the position of which depended on which visual area was being targeted for imaging. For V1 recordings, the craniotomy and head plate position was centred - 3.4 mm posterior and 2.8 mm lateral from bregma of the right hemisphere; for PM the position was slightly more medial, for AL and LM it was more lateral. The craniotomy was closed with a glass insert made from 3 layers of circular glass (#1 thickness; 1×5 mm, 2×3 mm diameter) bonded together with optical adhesive (Norland Products; catalogue no. 7106). The window was placed such that the 3 mm circular glass was in contact with the brain surface and the 5 mm diameter glass rested on the skull surrounding the craniotomy. The window was then sealed with dental cement. After surgery, all animals were allowed at least 1 week to recover before being imaged.

**Figure 1:**
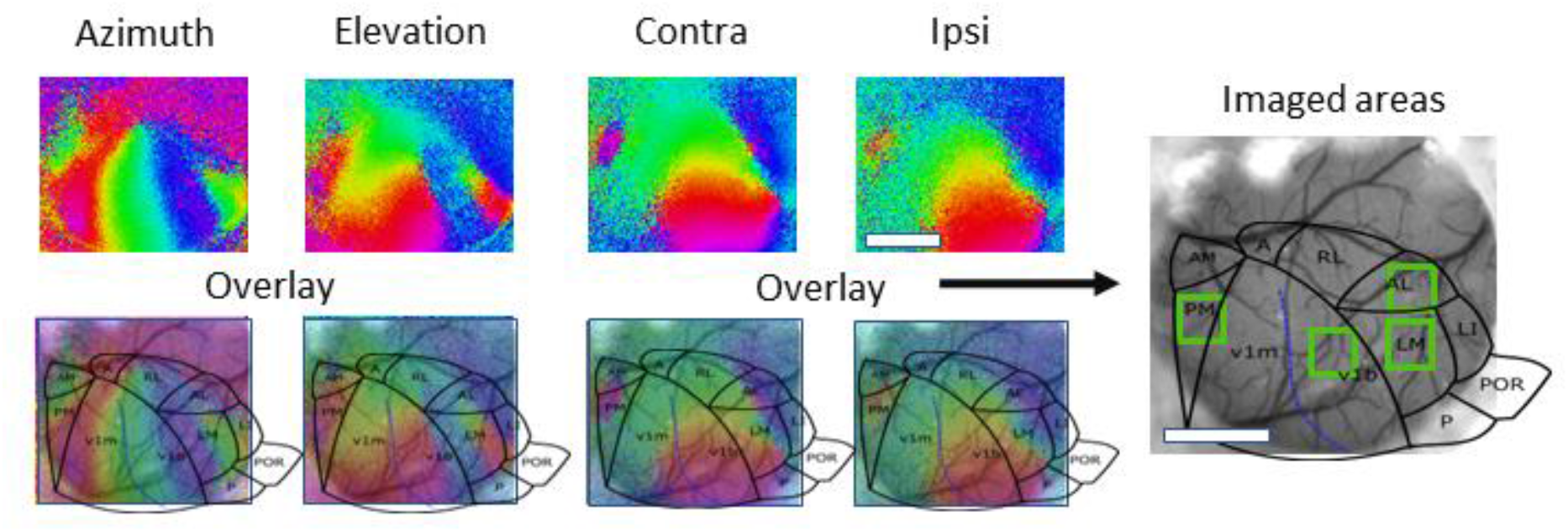
Identification of V1b, LM, AL and PM based on retinotopic mapping of the mouse visual cortex.

### Intrinsic Signal Imaging

Mice previously implanted with a cranial window and head-plate were placed under anaesthesia using isoflurane (5%) and oxygen (0.2 L/min). They were transferred to a heating pad to maintain body temperature at 37°C, and the head-plate was attached to a custom-made holder. Isoflurane was subsequently reduced to 1% for maintenance of anaesthesia throughout the imaging session.

The cortex was illuminated using a halogen light source with interchangeable bandpass filters. Initially green light (546nm) was used to acquire an image of the cortical surface vasculature. Intrinsic signal imaging (Imager 3001, Optical Imaging Inc, Israel) was carried out using red light (700 nm). The focal point of the camera was adjusted to 150-200 μm below the surface of the cortex. A screen was positioned 30 cm away from the nose of the mouse such that stimuli could be presented in the binocular and the left monocular visual field of the mouse (contralateral to the imaged right hemisphere).

To identify V1 and higher visual areas, a periodic stimulus protocol was used (Kalatsky and Stryker, 2003). Computer controlled shutters were used to present stimuli to one eye at a time. These were generated by VSG5 (Cambridge Research Systems) and consisted of a bar of 40 deg in length and 4 deg width drifting temporally upwards at a rate of 0.125 Hz and a speed of 15°/s, presented on a cathode ray tube screen positioned in front of the mouse at a distance of 14 cm. The stimulus was presented to each eye six times. The cortex was illuminated with red light (700±10 nm interference filter). Images were acquired with an Imager 3001 (Optical Imaging Inc, Mountainside, NJ) using an Adimec 50-R64 camera and running VDAQ software. For each pixel in the imaged region, phase and amplitude of the optical signal at 0.125 Hz were calculated by Fast Fourier Transform (Kalatsky and Stryker, 2003). Response amplitudes represent ΔR/R values (where R is light reflected), and phases represent retinotopic positions of maximal response (which is colour coded). The binocular zone of V1 was determined by thresholding the ipsilateral eye response map at 60% of the maximum pixel value.

### Two-photon imaging

In vivo 2-photon imaging was performed using a resonant scanning microscope (Thorlabs, B-Scope) with a 16x 0.8NA water-immersion objective (Nikon). GCaMP6f was excited at 980nm using a Ti:sapphire laser (Chameleon, Coherent) with a maximum laser power at sample of 50mW. Data were acquired at approximately 60Hz and averaged, resulting in a frame rate of approximately 10Hz. For all imaging experiments animals were head-fixed and placed on a custom designed cylindrical treadmill whose axis was fixed. Imaging and visual stimulation timing data were acquired using custom written DAQ code (Matlab) and a DAQ card (NI PCIe-6323, National Instruments). Visual stimuli were generated in Matlab using the psychophysics toolbox (Brainard, 1997), and displayed on a calibrated LCD screen (Iiyama, B2080HS; width x height 26 × 47 cm) placed 20 cm from the eyes in the binocular visual field of the animal. All visual stimuli were circular, unidirectionally drifting square wave gratings of uniform size (40×40 deg), spatial frequency (0.3 cycles per degree) and temporal frequency (1.25 Hz) presented pseudo randomly at 4 orientations of 0°, 45°, 90° and 135° degrees and at full contrast. Drifting gratings were presented for two seconds followed by a five second inter-trial interval during which a grey screen was displayed. Stimuli were shown to one eye at a time using computer-controlled shutters. In order to determine ocular dominance, stimuli were repeated 10 times in a pseudo-random order.

Recordings were made from a 400×400μm field of view, with the position for each HVA chosen on the basis of the retinotopic maps obtained with intrinsic signal imaging and comparison with published area maps of the mouse visual cortex (Wang and Burkhalter, 2007; Marshel et al., 2011); an example is shown in Fig. 1. The field of view was identical between recording sessions for each animal, and the region of interest chosen for each HVA was fixed within that field of view. Within the field of view all identifiable neurons were analysed, but they were not tracked individually across recording sessions.

Drifting vertical and horizontal bars were shown separately to the left (contralateral) and right (ipsilateral) eye using a periodic stimulation protocol (for details, see Methods). The stimulus phases producing the maximal response are colour coded and represent retinotopic azimuth and elevation, respectively. Each visual cortical area contains a retinotopic map, with boundaries between areas characterised by colour reversals. Individual areas were identified by overlaying a published map of higher visual areas in the mouse (Marshel et al., 2011). Scale bars, 1 mm.

### Calcium imaging data analysis

Calcium imaging data was registered and segregated into neuronal regions of interest using Suite2P (Pachitariu et al., 2017). Pixel-wise stimulus preference maps were constructed by first calculating the mean of the registered imaging frames recorded during the drifting phase of each stimulus, and then determining for each pixel the stimulus which elicited the largest mean response.

### Quantification and statistical analysis of visual responses

The visual responses of individual neurons were quantified as the mean dF/F value between 0.5-1.5 secs after stimulus onset, with a baseline period quantified for each trial as the mean dF/F between 1-0 secs before stimulus onset. Orientation tuning curves were fit to a 1d Gaussian function, constrained to peak at the preferred orientation, and with amplitude determined by the response amplitude at this orientation. Fitting was performed using the *fit* function in Matlab. Orientation tuning fit curves were used to calculate an orientation selectivity index OSI defined as OSI = 1 - circular variance, with circular variance calculated by the method of (Batschelet, 1981)

An ocular dominance index ODI was calculated for each neuron as 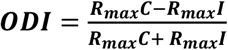

A neuron was classified as visually responsive if it showed dF/F during visual stimulation being significantly greater (at p < 0.01) for at least one stimulus than for baseline, with statistical significance calculated by Kruskal Wallis test between the baseline and visual stimulation periods. Kruskal Wallis tests to measure differences in response amplitude by stimulus orientation were used to assess whether neurons discriminated orientation for stimuli shown to each eye.

For each area of interest, significant changes in the proportions of recorded cells which have given properties over the course of the experiment were tested for by use of a continuity-corrected version of Wilson’s test of equal proportions (Wilson, 1927) using the prop.test function of the stat package in R. The properties of interest included the proportion of all cells being visually responsive, the proportion of visually responsive being contralateral, ipsilateral or binocularly responsive, the proportion of contralateral responsive cells being orientation selective, and the proportion of ipsilateral responsive cells being orientation selective.

Changes in ODI values of visually responsive cells over the course of the experiment, per area of interest were detected by use of Kruskal Wallis tests. In areas where there were found to be significant differences in ODI across sessions, Wilcoxon tests were then carried out to test for significant differences in ODI between Baseline and immediately-post Monocular Deprivation sessions, and then between the post-Monocular deprivation and 2 weeks binocular recovery sessions. Changes in OSI for contralateral responsive cells, and for ipsilateral responsive cells were tested for in the same manner.

Correction for multiple testing was implemented by use of the conservative Bonferroni method, to minimize our reporting of false-positive results. Where adjusted p values failed to reach significance but the unadjusted values did we report both values, such that potential false-negative results likely to arise from stringent multiple comparison correction may also be identified.

## Results

We obtained complete V1b data sets consisting of pre-MD baseline, immediately post-MD and 7d binocular recovery time points, from 10 mice, LM data sets from 8 mice, AL recordings from 12 and PM recordings from 7 animals. We were able to further record 14d binocular recovery data from 7 animals for V1b, 6 for LM, 10 for AL and 6 for PM. The fraction of visually responsive neurons as well as the classification of the visually responsive neurons as contralaterally, binocularly or ipsilaterally responding is shown in Fig. 2. In the baseline condition between a third and half of neurons in V1b, LM and AL were visually responsive according to our criteria (41.5%, 46.9% and 36.2%, respectively), but only 13.8% in PM – our visual stimulus parameters were possibly not optimal for this area. The percentage of visually responsive neurons varied over time significantly in all areas (Fig. 2A), as a result of monocular deprivation and binocular recovery; however, changes were not significant for PM once adjustment for multiple comparisons was made (Wilson test for equality of proportions; V1b: p=2.06×10^−54^, *χ*^2^ = 260.2; LM: p=2.88×10^−23^, p=5.75×10^−25^, *χ*^2^ = 116.0; AL: p=1.56×10^−19^, p=3.11×10^−21^, *χ*^2^ = 98.6; PM: adjusted p=0.118, unadjusted p=0.00236, *χ*^2^ = 14.44.)

**Figure 2:**
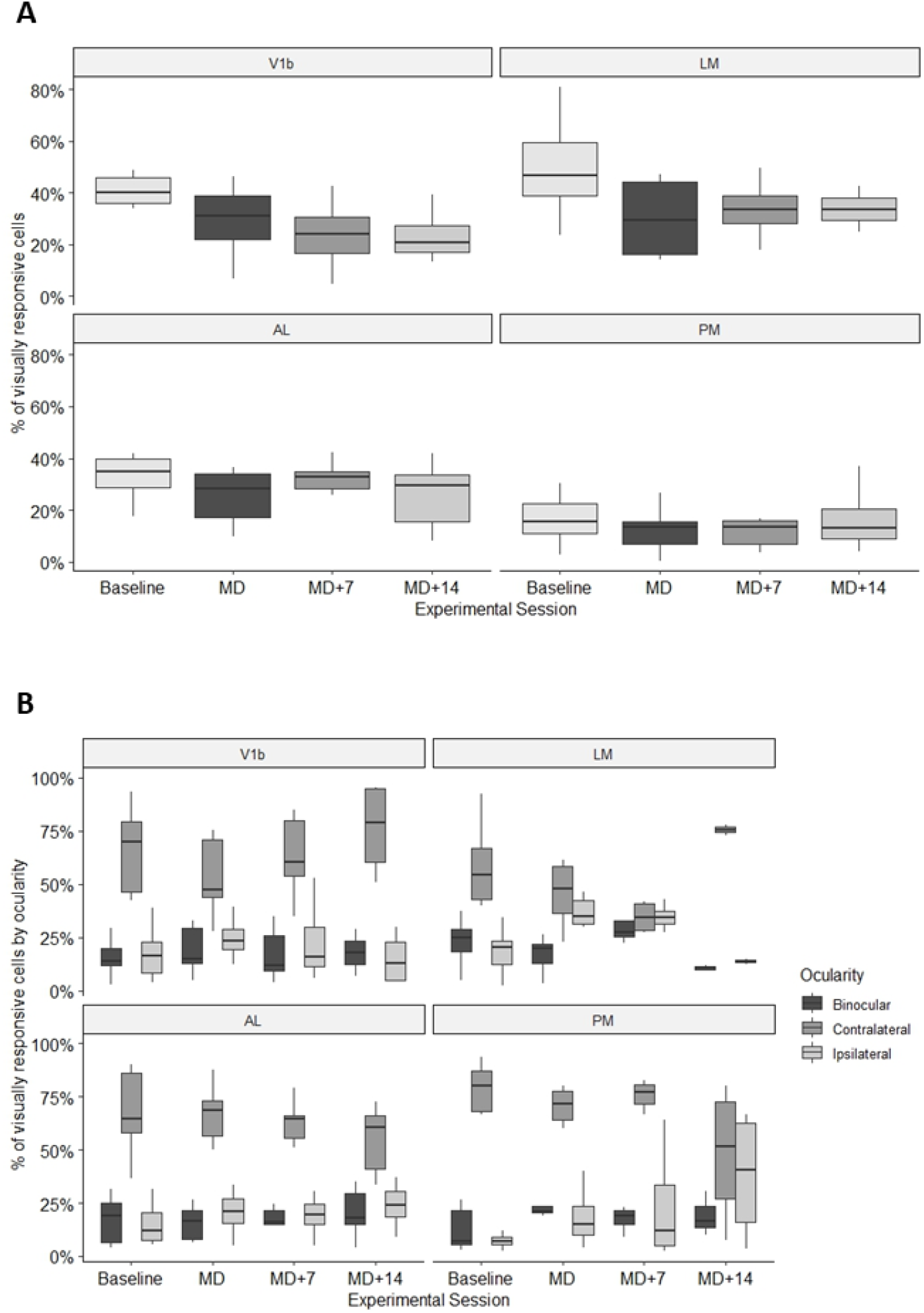
Proportions of visually responsive cells across visual areas and timepoints. A) Percentage of cells significantly responsive to visual stimulation out of all cells identified with V1b, LM, AL and PM across time points. Box plots showing medians and interquartile ranges as well as minimum and maximum values. B) Proportion (out of all visually responsive cells) of identified neurons in V1b, LM, AL and PM that responded either through the contralateral eye only (dark grey bars), through both eyes (mid grey) or through the ipsilateral eye only (light grey). Box plots show medians and interquartile ranges as well as minimum and maximum values.

### Ocular dominance

V1b, AL and LM contained 16-22% of binocularly responsive cells (as a proportion of all visually responsive cells; V1b: 16.8%, LM 22.1%, AL: 16.6%), but PM only 13.6% (Fig. 2B). While in V1b and LM around 60% of the visually responsive neurons responded exclusively through the contralateral eye (V1b: 62.4%, LM: 59.8%), for the dorsal stream areas AL and PM that figure was considerably higher (AL: 68.3%, PM: 78.8%). The proportion of cells responding exclusively through the ipsilateral eye in the baseline condition was highest in V1b (28.7%), and at a similar level in all 3 higher visual areas (LM: 21.1%; AL: 22.6%; PM: 23.4%).

The percentage of visually responsive cells which were responsive to the contralateral eye only changed significantly over the course of monocular deprivation (of that eye) and binocular recovery in all areas; however, changes were not significant for PM once adjustment for multiple comparisons was made (Wilson test for equality of proportions; V1b: p= 8.40×10^−10^, *χ*^2^ = 53.17; LM: p=6.60×10^−9^, *χ*^2^ = 48.98; AL: p=2.77×10^−9^, *χ*^2^=50.75; PM: adjusted p=0.162, unadjusted p=0.00323, *χ*^2^ =13.77).

Two weeks of MD caused a decrease in the proportion of neurons responding exclusively through the contralateral (deprived) eye in all 4 cortical areas; this reduction was greatest in V1b (13.7%), and somewhat smaller in LM (7.58%) AL (6.53%) and PM (11.5%).

There were striking differences between the four cortical areas in the proportion of exclusively contralateral neurons during the binocular recovery period following reopening of the deprived (contralateral) eye. This proportion returned to pre-MD levels within a week in V1b, and continued to grow in the second week, while it decreased further for a week and then more than recovered in ventral-stream area LM (Fig. 2 B). In contrast, it continued to fall in the dorsal-stream areas AL and PM (Fig. 2 B).

In areas V1b and LM (but not the dorsal-stream areas AL and PM), the percentage of visually responsive cells which were binocularly responsive changed significantly with experience of monocular deprivation and binocular recovery (V1b: p=0.0384, *χ*^2^ = 16.82; LM: p=0.00152, *χ*^2^ = 23.59; AL: adjusted p>0.99, unadjusted p=0.0305, *χ*^2^ = 8.909; PM: unadjusted p = 0.394, *χ*^2^= 2.980).

In V1b, the proportion of binocularly responsive neurons over time mirrors the percentage of contralaterally responsive cells, increasing after 2 weeks of MD and then returning to pre-MD levels within a week of binocular recovery. In contrast, the percentage of binocularly responsive neurons in LM increased one week after reopening of the deprived eye but then decreased to well below baseline levels after two weeks. In the dorsal stream areas AL and PM the percentage of binocularly responsive neurons changed little over the course of MD and binocular recovery.

In V1b, the proportion of cells responsive only to the ipsilateral eye did not change significantly over the course of the experiment (V1b: p>0.99, *χ*^2^ =1.5). In all 3 of the higher visual areas LM, AL and PM, the percentage of cells which were responsive only to ipsilateral eye stimulation changed over time; however, changes were not significant for PM once adjustment for multiple comparisons was made (LM: p=3.56×10^−18^, *χ*^2^ =92.3; AL: p=1.33 ×10^−4^, *χ*^2^ =28.6; PM: adjusted p=0.645, unadjusted p=0.0129, *χ*^2^ =10.8).

In area LM the reduction in contralaterally responsive neurons during MD was mirrored by a large increase in the proportion of ipsilaterally responsive cells, from 21.1% to 33.9%. In V1b, the proportion of exclusively ipsilaterally responsive cells stayed stable during the MD period while in AL and PM it increased slightly during MD and continued to increase during 2 weeks of binocular recovery (Fig. 2B).

Effects of monocular deprivation and recovery after reopening of the deprived eye are more commonly assessed through the ocular dominance index (ODI) although this measure does not reveal the underlying changes in response through the individual eyes. Overall, ODI changed over the course of the experiment in all 4 areas (Kruskal-Wallis, V1b: p=4.60×10^−4^, *χ*^2^ =26.1; LM: p=6.10×10^−9^, *χ*^2^ =49.1; AL: p=3.71×10^−5^, *χ*^2^ =31.3; PM: p=0.0090, *χ*^2^ =19.9). An example of the changes seen for all visually responsive cells in area AL of one animal over time is shown in Fig. 3. Population data for all areas across time are shown in Fig. 4.

**Figure 3:**
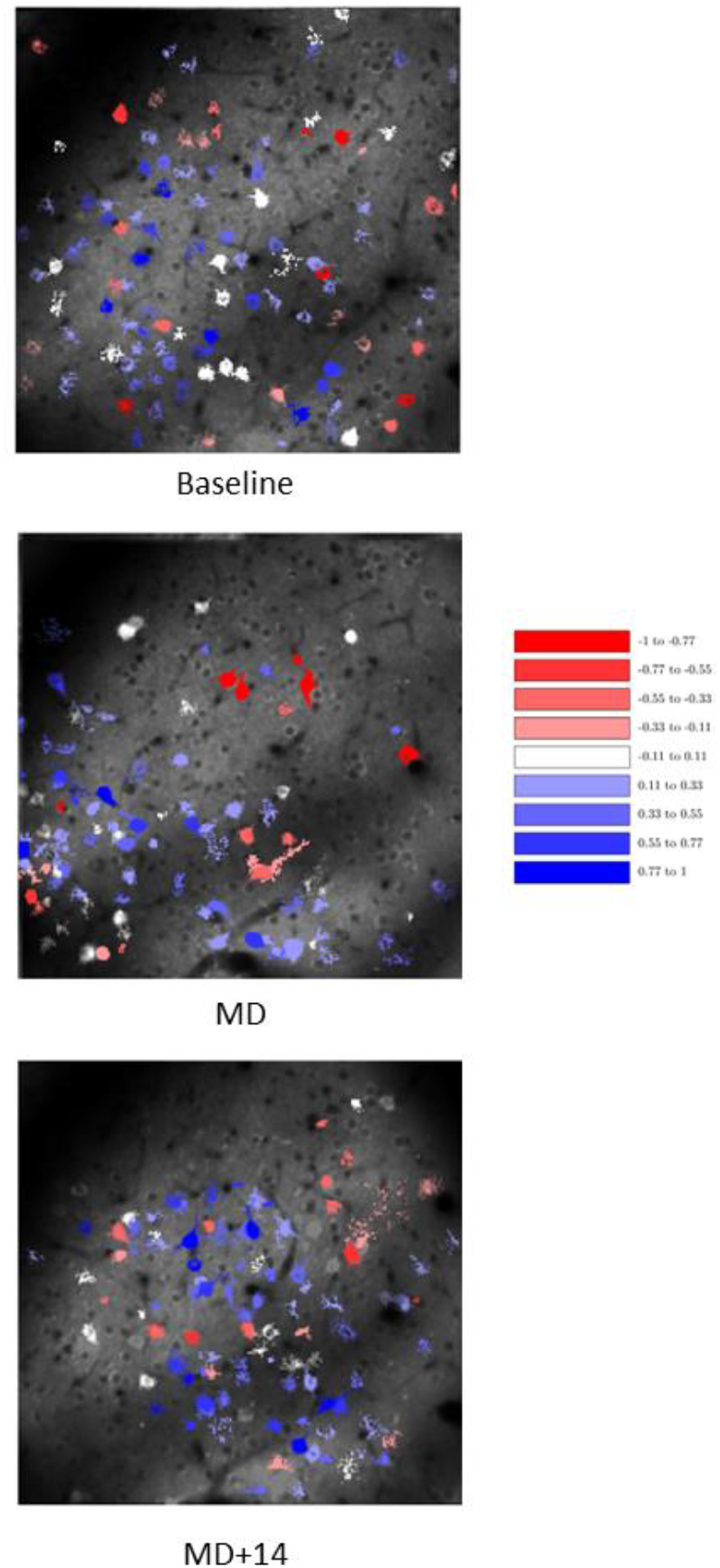
Changes to the ocular dominance index over time of all visually responsive cells in area AL of one mouse.

**Figure 4:**
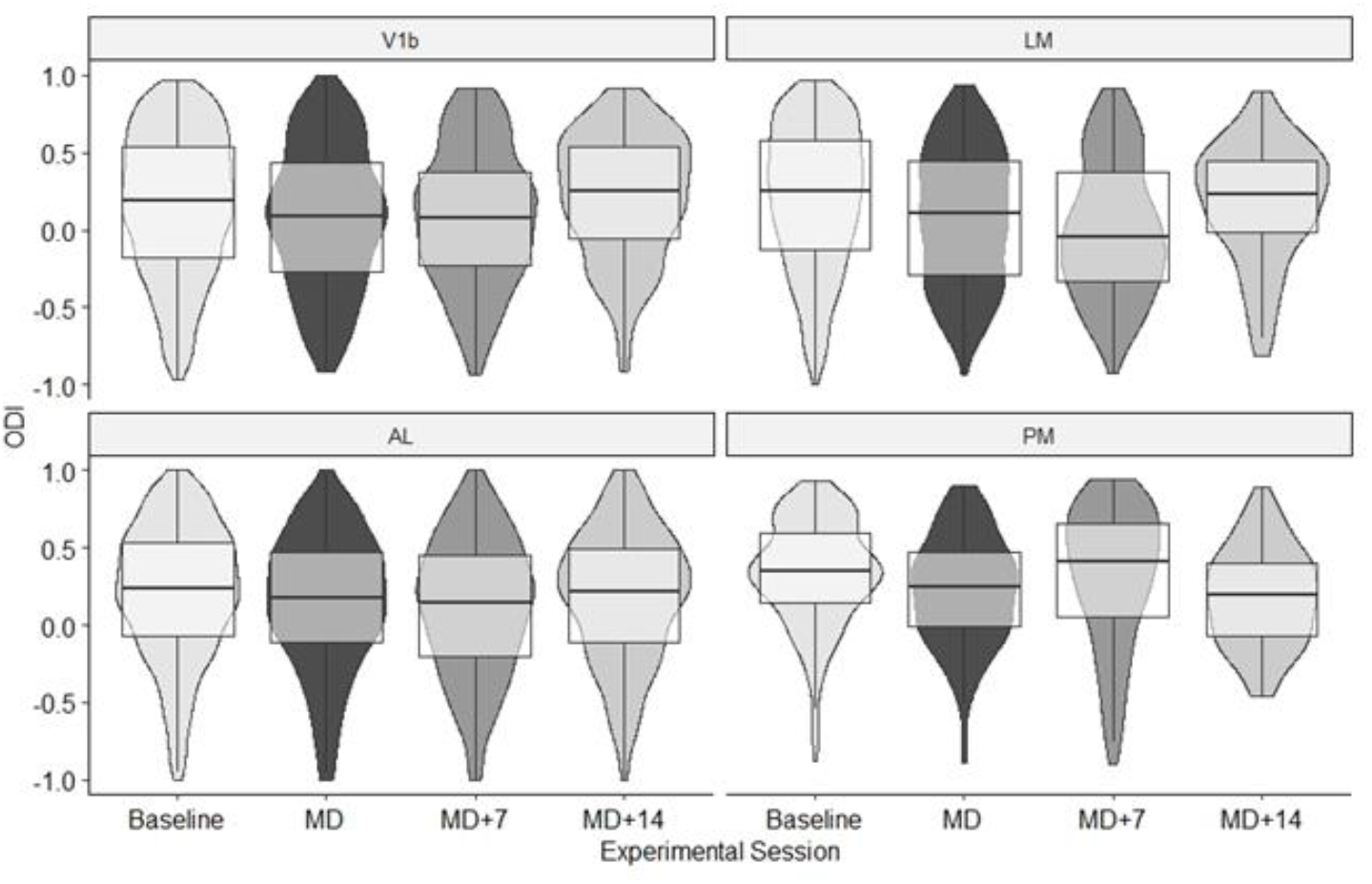
Changes to ocular dominance index caused by monocular deprivation and binocular recovery.

All visually responsive cells in the baseline recording (top) were colour coded after dividing the population into 9 equal bands ranging from ODI = -1.0 (red) to ODI = 1.0 (blue). The same ODI bands were then used to colour code the visually responsive cells following MD (middle) and after 14 days of binocular recovery (bottom).

Ocular dominance index (ODI) for visual areas V1b, LM, AL and PM across timepoints. Violin plots show medians and interquartile ranges, minimum and maximum values as well as density of ODI values of individual neurons. Note that these have been normalised for the overall number of neurons imaged for each timepoint and cortical area.

As expected, in V1b there was a significant shift in ODI from 0.195 to 0.084 during 2 weeks of MD (Wilcoxon, p=0.00427, W = 921661). During 2 weeks of recovery there was a significant shift back to an ODI of 0.256 (Wilcoxon, p=0.0172, *χ*^2^ = 46517). In comparison, in the ventral-stream area LM we observed a more pronounced shift in ODI from 0.248 to 0.105 (Wilcoxon, p=2.21×10^−4^, W = 350405), and a nearly full recovery to 0.235 during the subsequent 2 weeks of binocular vision, although this did not quite reach significance (Wilcoxon, unadjusted p=0.0554, W= 14482).

In AL the decrease in ODI (from 0.237 to 0.179) during 2 weeks of MD was more modest but still significant after correction for multiple comparison (Wilcoxon, p=0.0266, W = 826338); the ODI then decreased further to 0.149 after one week of binocular vision before returning to near baseline after two weeks (Wilcoxon, p=0.148, W=235673). In PM, the ODI also decreased during MD, from 0.351 to 0.251, but this was not quite significant after correction for multiple comparison (Wilcoxon, adjusted p= 0.0713, unadjusted p=0.00143, W = 27596). After one week of recovery ODI increased to 0.411 but then decreased again, to 0.193 after 2 weeks of binocular vision; this shift was not significant (Wilcoxon, unadjusted p=0.455, *χ*^2^ = 3875).

In summary, a very strong and significant shift in ODI was seen in the ventral steam area LM following MD; the OD shift in V1b was smaller, and those in dorsal-steam areas AL and PM were smaller still. Within 2 weeks of restored binocular vision, a recovery of the ODI was observed in V1b, LM and AL, but not in PM (Fig. 4).

### Orientation selectivity

It is worth noting that the percentage of orientation selective cells out of all cells responding through the contralateral eye was above 50% not only in area V1b (51.0%), but also in LM (63.2%) and AL (54.3%), while it was only 22.4% in area PM (Fig. 5A). Following MD and recovery we observed significant changes in the orientation selectivity index (OSI) of those orientation selective cells responsive only to contralateral eye stimulation in area AL (Kruskal-Wallis: p= 2.10×10^−5^, *χ*^2^ = 32.5) and LM (KW, p= 4.31×10^−5^, *χ*^2^ = 31.0); changes in V1 fell just short of significance after correction for multiple comparisons (KW: adjusted p= 0.0635, unadjusted p=0.00127, *χ*^2^ = 15.8). There were no significant changes in OSI in PM (KW, unadjusted p=0.654, *χ*^2^ = 1.62)

**Figure 5:**
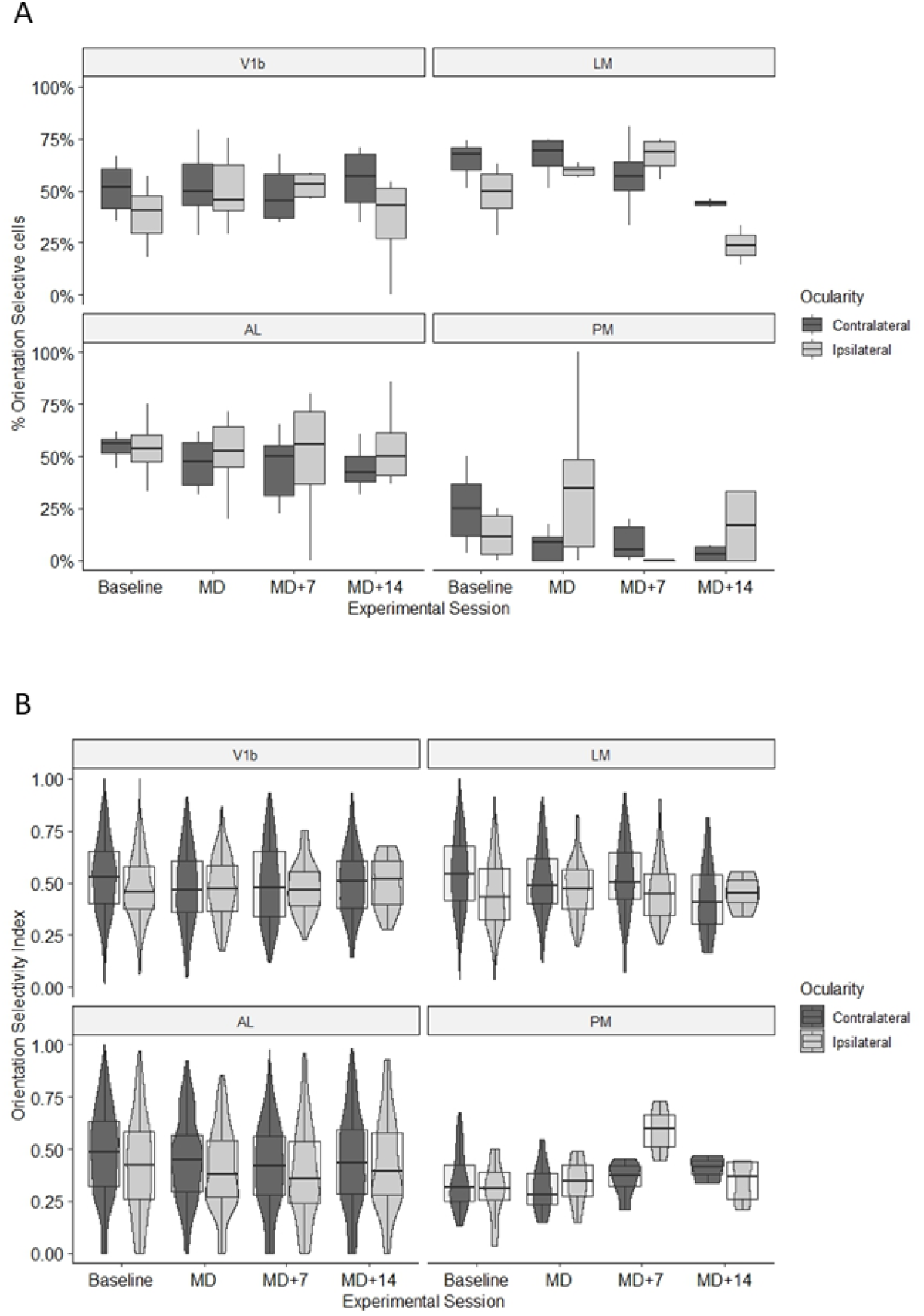
Effects of monocular deprivation binocular recovery on orientation selectivity of visually responsive neurons in cortical areas V1b, LM, AL and PM. A) Percentage of orientation selective cells responding to contralateral and ipsilateral eye stimulation, respectively, before and after monocular deprivation and following binocular recovery. Box plots showing medians and interquartile ranges as well as minimum and maximum values. B) Orientation selectivity index (OSI) of cells responding to contralateral and ipsilateral eye stimulation, respectively, before and after monocular deprivation and following binocular recovery. Box plots showing medians and interquartile ranges as well as minimum and maximum values

The orientation selectivity index of responses through the contralateral (deprived) eye was reduced following monocular deprivation in V1b, from a median OSI before MD of 0.528 and a median OSI after 2 weeks of MD of 0.475, but this reduction failed to reach significance after correction for multiple comparison (Wilcoxon test, adjusted p= 0.0725, unadjusted p = 0.00145, W = 75387). During 2 weeks of binocular vision there was a nearly complete recovery to a median OSI of 0.521 (Fig. 5B).

In ventral-stream area LM we also observed a decrease in OSI of cells responding through the contralateral eye, from a median 0.551 before MD to 0.485 immediately after MD. Again, this decrease did not reach significance after correcting for multiple comparisons (Wilcoxon test, adjusted p=0.0886, unadjusted p=00177, W = 46422). There was a partial recovery after one week of binocular vision to a median OSI of 0.502; there were not enough cells meeting our criteria for an evaluation after two weeks of recovery.

In the dorsal-stream area AL the orientation selectivity index of responses through the contralateral eye decreased from a median baseline of 0.486 to 0.440 after 2 weeks of MD; however, this was not significant (Wilcoxon test, adjusted p = 0.758, unadjusted p = 0.015, W = 87627). Surprisingly, the OSI then decreased further to 0.402 after 2 weeks of binocular vision.

Orientation selectivity of responses through the contralateral eye was substantially lower in PM than in V1b, LM or AL, not only in terms of the proportion of orientation selective cells but also their OSI; this was unaffected by MD, with a median OSI before MD of 0.321 and a median OSI after MD of 0.356.

Orientation selectivity of responses through the ipsilateral (non-deprived) eye did not change significantly over time in any of the areas tested (Kruskal-Wallis, p>0.99, for all 4 areas).

## Discussion

The observed effects of monocular deprivation on visual responses in V1b confirm results previously reported: MD during the critical period causes a decrease in responsiveness of neurons to stimulation of the deprived eye (Gordon and Stryker, 1996; Frenkel and Bear, 2004) due to long-term depression (Yoon et al., 2009), and it causes a reduction in orientation selectivity of those deprived-eye responses. Both the amplitude of responses and their orientation selectivity largely recover during a subsequent period of concordant binocular vision. The overall pattern of effects of MD and recovery was similar in area LM (considered part of the ventral stream), as was the decrease in orientation selectivity through the deprived eye. In contrast, area AL and area PM (in the dorsal stream) exhibited smaller changes in the response through the deprived eye, the overall ocular dominance, and the orientation selectivity of responses.

There are surprisingly few reports on experience dependent, and more specifically ocular dominance, plasticity in higher visual cortical areas from any mammalian species. Ocular dominance plasticity is widely viewed as a consequence of the convergence of retinal inputs representing the two eyes and a prerequisite for the precise matching of the tuning of response to stimulation of the two eyes (Wang et al., 2010; Gu and Cang, 2016; Chang et al., 2020; Tan et al., 2020). If this is the case, then one might expect OD plasticity to occur principally in neurons in the primary visual cortex since eye-of-origin information is largely lost beyond it in all species studied so far, and to be subsequently ‘inherited’ by the higher visual areas that receive input from V1. If on the other hand ocular dominance plasticity is linked to the development of disparity sensitivity, then one might expect to observe it in higher visual areas where such selectivity is present. Outside of V1, disparity tuning of neurons has been reported for areas LM and RL, which are the areas with the largest representation of the binocular visual field in mice (Garrett et al., 2014; Zhuang et al., 2017). Area RL is tuned mainly to near disparities, representing visual stimuli very close to the mouse (La Chioma et al., 2019), while area LM (but not RL) exhibits true stereo sensitivity, with neurons responding more strongly to correlated than to anti-correlated random-dot correlograms (La Chioma et al., 2020).

Such responses are typical of visual areas of the ventral stream in primates, namely V4 and inferotemporal cortex. It is therefore interesting to note that LM exhibited a significant OD shift in response to monocular deprivation, of greater magnitude than that observed in V1. This finding is paralleled by similar results from cat area 21a, considered the gateway to the ventral stream in that species, where OD plasticity has also been found to be stronger (and more long-lasting) than in V1 (Wang et al., 2019).

The fact that the ocular dominance shift following MD is greater in ventral-stream area LM than in V1b (while it is smaller in the dorsal stream areas AL and PM) suggests that changes in ocular dominance in higher visual areas are not simply inherited from V1b. For primates, it has long been known that the severity of amblyopia as assessed by tests of visual acuity cannot be (fully) explained by the neuronal response deficits in the primary visual cortex (Kiorpes et al., 1998; Kiorpes and McKee, 1999), suggesting that responses in higher visual areas are more strongly affected. Similarly, in cats reopening of a previously deprived eye leads to only a partial recovery of visual acuity in that eye despite a full restoration of balanced ocular dominance and normal orientation selectivity (Kind et al., 2002), indicating persisting deficits in higher visual areas. The relationship between basic visual functions such as visual acuity and neuronal response properties in higher visual cortical areas requires further study across species.

## CRediT authorship contribution statement

Rosie Craddock: Methodology, Investigation, Software, Formal analysis, Writing - review & editing, Visualization; Asta Vasalauskaite; Methodology, Investigation, Formal analysis, Writing - review & editing; Adam Ranson: Conceptualization, Methodology, Writing - review & editing; Frank Sengpiel: Conceptualization, Methodology, Project administration, Writing - original draft, Writing - review & editing, Visualization, Funding acquisition

## Acknowledgements

We thank Fangli Chen for expert technical assistance.

## References

Barkus C, Bergmann C, Branco T, Carandini M, Chadderton PT, Galiñanes GL, Gilmour G, Huber D, Huxter JR, Khan AG, King AJ, Maravall M, O’Mahony T, Ragan CI, Robinson ESJ, Schaefer AT, Schultz SR, Sengpiel F, Prescott MJ (2022) Refinements to rodent head fixation and fluid/food control for neuroscience. Journal of Neuroscience Methods:109705.

Batschelet E (1981) Circular Statistics in Biology. London: Academic Press.

Bear MF, Kleinschmidt A, Gu QA, Singer W (1990) Disruption of experience-dependent synaptic modifications in striate cortex by infusion of an NMDA receptor antagonist. Journal of Neuroscience 10:909–925.

Brainard DH (1997) The Psychophysics Toolbox. Spat Vis 10:433–436.

Chang JT, Whitney D, Fitzpatrick D (2020) Experience-Dependent Reorganization Drives Development of a Binocularly Unified Cortical Representation of Orientation. Neuron 107:338-350.e335.

Dews PB, Wiesel TN (1970) Consequences of monocular deprivation on visual behaviour in kittens. J Physiol 206:437–455.

Dräger UC (1975) Receptive fields of single cells and topography in mouse visual cortex. Journal of Comparative Neurology 160:269–289.

Dräger UC (1978) Observations on monocular deprivation in mice. Journal of Neurophysiology 41:28–42.

Espinosa JS, Stryker MP (2012) Development and Plasticity of the Primary Visual Cortex. Neuron 75:230–249.

Frenkel MY, Bear MF (2004) How Monocular Deprivation Shifts Ocular Dominance in Visual Cortex of Young Mice. Neuron 44:917–923.

Garrett ME, Nauhaus I, Marshel JH, Callaway EM (2014) Topography and Areal Organization of Mouse Visual Cortex. The Journal of Neuroscience 34:12587–12600.

Gordon JA, Stryker MP (1996) Experience-dependent plasticity of binocular responses in the primary visual cortex of the mouse. JNeurosci 16:3274–3286.

Greifzu F, Pielecka-Fortuna J, Kalogeraki E, Krempler K, Favaro PD, Schlüter OM, Löwel S (2014) Environmental enrichment extends ocular dominance plasticity into adulthood and protects from stroke-induced impairments of plasticity. Proceedings of the National Academy of Sciences 111:1150–1155.

Gu Y, Cang J (2016) Binocular matching of thalamocortical and intracortical circuits in the mouse visual cortex. eLife 5:e22032.

Harwerth RS, Smith EL, III, Boltz RL, Crawford MLJ, von Noorden GK (1983) Behavioral studies on the effect of abnormal early visual experience in monkeys: spatial modulation sensitivity. Vision Res 23:1501–1510.

Hubel DH, Wiesel TN (1970) The period of susceptibility to the physiological effects of unilateral eye closure in kittens. Journal of Physiology 206:419–436.

Hubel DH, Wiesel TN, LeVay S (1977) Plasticity of ocular dominance columns in monkey striate cortex. PhilTransRSocLondB 278:377–409.

Kalatsky VA, Stryker MP (2003) New paradigm for optical imaging: temporally encoded maps of intrinsic signal. Neuron 38:529–545.

Kaneko M, Stellwagen D, Malenka RC, Stryker MP (2008a) Tumor Necrosis Factor-[alpha] Mediates One Component of Competitive, Experience-Dependent Plasticity in Developing Visual Cortex. Neuron 58:673–680.

Kaneko M, Hanover JL, England PM, Stryker MP (2008b) TrkB kinase is required for recovery, but not loss, of cortical responses following monocular deprivation. Nat Neurosci 11:497–504.

Kind PC, Mitchell DE, Ahmed B, Blakemore C, Bonhoeffer T, Sengpiel F (2002) Correlated binocular activity guides recovery from monocular deprivation. Nature 416:430–433.

Kiorpes L, McKee SP (1999) Neural mechanisms underlying amblyopia. CurrOpinNeurobiol 9:480–486.

Kiorpes L, Daw N (2018) Cortical correlates of amblyopia. Visual Neuroscience 35:E016.

Kiorpes L, Kiper DC, O’Keefe LP, Cavanaugh JR, Movshon JA (1998) Neuronal correlates of amblyopia in the visual cortex of macaque monkeys with experimental strabismus and anisometropia. Journal of Neuroscience 18:6411–6424.

La Chioma A, Bonhoeffer T, Hübener M (2019) Area-Specific Mapping of Binocular Disparity across Mouse Visual Cortex. Current Biology 29:2954-2960.e2955.

La Chioma A, Bonhoeffer T, Hübener M (2020) Disparity Sensitivity and Binocular Integration in Mouse Visual Cortex Areas. The Journal of Neuroscience 40:8883–8899.

Levelt CN, Hübener M (2012) Critical-Period Plasticity in the Visual Cortex. Annual Review of Neuroscience 35:309–330.

Marshel James H, Garrett Marina E, Nauhaus I, Callaway Edward M (2011) Functional Specialization of Seven Mouse Visual Cortical Areas. Neuron 72:1040–1054.

Mrsic-Flogel TD, Hofer SB, Ohki K, Reid RC, Bonhoeffer T, Hübener M (2007) Homeostatic Regulation of Eye-Specific Responses in Visual Cortex during Ocular Dominance Plasticity. Neuron 54:961–972.

Murakami T, Matsui T, Ohki K (2017) Functional Segregation and Development of Mouse Higher Visual Areas. The Journal of Neuroscience 37:9424–9437.

Olson CR, Freeman RD (1978) Monocular deprivation and recovery during sensitive period in kittens. J Neurophysiol 41:65–74.

Pachitariu M, Stringer C, Dipoppa M, Schröder S, Rossi LF, Dalgleish H, Carandini M, Harris KD (2017) Suite2p: beyond 10,000 neurons with standard two-photon microscopy. bioRxiv:061507.

Ranson A, Cheetham CE, Fox K, Sengpiel F (2012) Homeostatic plasticity mechanisms are required for juvenile, but not adult, ocular dominance plasticity. Proceedings of the National Academy of Sciences 109:1311–1316.

Sawtell NB, Frenkel MY, Philpot BD, Nakazawa K, Tonegawa S, Bear MF (2003) NMDA Receptor-Dependent Ocular Dominance Plasticity in Adult Visual Cortex. Neuron 38:977–985.

Tan L, Tring E, Ringach DL, Zipursky SL, Trachtenberg JT (2020) Vision Changes the Cellular Composition of Binocular Circuitry during the Critical Period. Neuron 108:735-747.e736.

Wang BS, Sarnaik R, Cang J (2010) Critical Period Plasticity Matches Binocular Orientation Preference in the Visual Cortex. Neuron 65:246–256.

Wang J, Ni Z, Jin A, Yu T, Yu H (2019) Ocular Dominance Plasticity of Areas 17 and 21a in the Cat. Frontiers in Neuroscience 13.

Wang Q, Burkhalter A (2007) Area map of mouse visual cortex. J Comp Neurol 502:339–357.

Wang Q, Gao E, Burkhalter A (2011) Gateways of Ventral and Dorsal Streams in Mouse Visual Cortex. The Journal of Neuroscience 31:1905–1918.

Wang Q, Sporns O, Burkhalter A (2012) Network Analysis of Corticocortical Connections Reveals Ventral and Dorsal Processing Streams in Mouse Visual Cortex. The Journal of Neuroscience 32:4386–4399.

Wiesel TN, Hubel DH (1963) Single-cell responses in striate cortex of kittens deprived of vision in one eye. JNeurophysiol 26:1003–1017.

Wilson EB (1927) Probable Inference, the Law of Succession, and Statistical Inference. Journal of the American Statistical Association 22:209–212.

Yoon BJ, Smith GB, Heynen AJ, Neve RL, Bear MF (2009) Essential role for a long-term depression mechanism in ocular dominance plasticity. Proceedings of the National Academy of Sciences 106:9860–9865.

Zhuang J, Ng L, Williams D, Valley M, Li Y, Garrett M, Waters J (2017) An extended retinotopic map of mouse cortex. eLife 6:e18372.

